# Thermodynamic rigidity of harmonic brain states relates to general mental ability in juvenile myoclonic epilepsy

**DOI:** 10.64898/2026.04.06.715875

**Authors:** Felipe Branco De Paiva, Meixian Zhao, Meishu Zhao, Santiago Philibert-Rosas, Cameron J. Brace, Erika Moe, Steven E. Haworth, Bruce P. Hermann, Moo K. Chung, Aaron F. Struck

## Abstract

Cognitive difficulties are increasingly recognized in juvenile myoclonic epilepsy (JME), but scalable biomarkers linking resting-state brain dynamics to general mental ability remain limited. Here, we combined topological data analysis, graph signal processing, machine learning, inverse Langevin modeling, and biophysical simulations to test whether EEG-derived network dynamics capture individual differences in general mental ability in JME.

We studied 54 patients with JME and 45 healthy controls using resting-state high-density EEG and the raw estimated full-scale score derived from the Wechsler Abbreviated Scale of Intelligence (WASI), used here as an index of general mental ability. Subject-specific low-alpha activity was reconstructed with generalized eigendecomposition, and graph-derived features were extracted from the projection of topological and alpha-power signals onto the functional connectome, providing a graph-harmonic description of large-scale brain-state dynamics. In controls, dynamic EEG-derived features significantly predicted general mental ability, whereas the same framework failed in JME. Because prediction in controls was driven mainly by dynamic measures of smoothness (Dirichlet energy), we next examined the temporal organization of alpha-power smoothness using an inverse Langevin framework. Within the patient group, greater thermodynamic rigidity—that is, stronger confinement of fluctuations around preferred network states—was associated with lower general mental ability. Relative to controls, patients also showed lower thermodynamic noise, indicating a reduced tendency to explore alternative network regimes.

Biophysical simulations suggested that reduced dendritic arborization can generate rigidity directly, whereas pharmacological stabilization of hyperexcitable circuits can shift the system toward a more rigid, lower-noise regime. Together, these findings suggest that cognition in JME is linked not only to altered resting-state network dynamics but also to stronger confinement of network-state fluctuations, with both intrinsic circuit abnormalities and treatment-related stabilization representing plausible routes to this rigid phenotype.

## Introduction

Epilepsy is among the most prevalent chronic neurological disorders worldwide, affecting more than 50 million individuals and contributing substantially to disability, health care utilization, and psychosocial burden (World Health Organization, 2024). While defined clinically by recurrent unprovoked seizures, epilepsy is increasingly understood as a disorder of large-scale brain networks, with consequences that extend beyond ictal events, including cognition (Bernhardt et al., 2015; Royer et al., 2022).

Juvenile myoclonic epilepsy (JME) accounts for approximately 5%–10% of epilepsies and has traditionally been defined by its characteristic seizure semiology, including bilateral myoclonic jerks and frequent generalized tonic–clonic seizures (Baykan & Wolf, 2017). However, JME is increasingly recognized as a broader disorder involving frontal-thalamocortical dysfunction, with convergent neuroimaging and neuropsychological evidence showing abnormalities that extend beyond seizure generation alone (Anderson & Hamandi, 2011; Wandschneider et al., 2012; Zhang et al., 2022; Garcia-Ramos et al., 2025). Cognitive difficulties in JME are particularly evident in executive and related frontal-lobe functions, and they can persist even in otherwise clinically stable patients (Wandschneider et al., 2012; Chawla et al., 2021; Struck et al., 2025). Because these difficulties emerge during adolescence and early adulthood, they may affect educational, occupational, and social trajectories. Consistent with this, JME has been associated with worse academic and less favorable long-term social outcomes in at least a subset of patients (Struck et al., 2025; Schneider-von Podewils et al., 2014).

Early identification of cognitive vulnerability in epilepsy remains difficult. Comprehensive neuropsychological assessment is clinically valuable, but it is resource-intensive and not consistently integrated into routine epilepsy care (Wilson et al., 2015). Interpretation of cognitive difficulties is also complicated by the fact that cognition may reflect multiple interacting influences, including syndrome-related network dysfunction, seizure burden, and antiseizure medication effects (Wilson et al., 2015; Eddy et al., 2011; Reyes et al., 2026). These limitations create a need for scalable biomarkers that can identify cognitive risk from data already acquired during standard clinical evaluation, particularly EEG (Wilson et al., 2015).

General mental ability, indexed here by the raw estimated full-scale score from the Wechsler Abbreviated Scale of Intelligence (WASI), is a useful outcome for this purpose because it captures variance shared across diverse cognitive tasks and predicts important life outcomes, including educational attainment, occupational functioning, health, and longevity (Deary et al., 2010; Deary, 2012). Resting-state EEG is an attractive candidate source of such biomarkers because it is already widely collected in epilepsy care, yet its value for capturing cognition-relevant large-scale dynamics remains underdeveloped.

Here we combined topological data analysis and graph signal processing to characterize how subject-specific low-alpha EEG activity is organized on functional networks, and we first tested whether these descriptors could predict individual differences in general mental ability (Fig. 1). Because prediction succeeded in controls but failed in JME, and because the control model was driven mainly by dynamic smoothness features, we next asked whether the temporal organization of smoothness itself differed in a cognitively meaningful way. Finally, to probe possible circuit-level origins of the observed dynamical patterns, we used biophysical simulations to test whether candidate microcircuit abnormalities and pharmacological stabilization could generate similar regimes.

**Figure 1:**
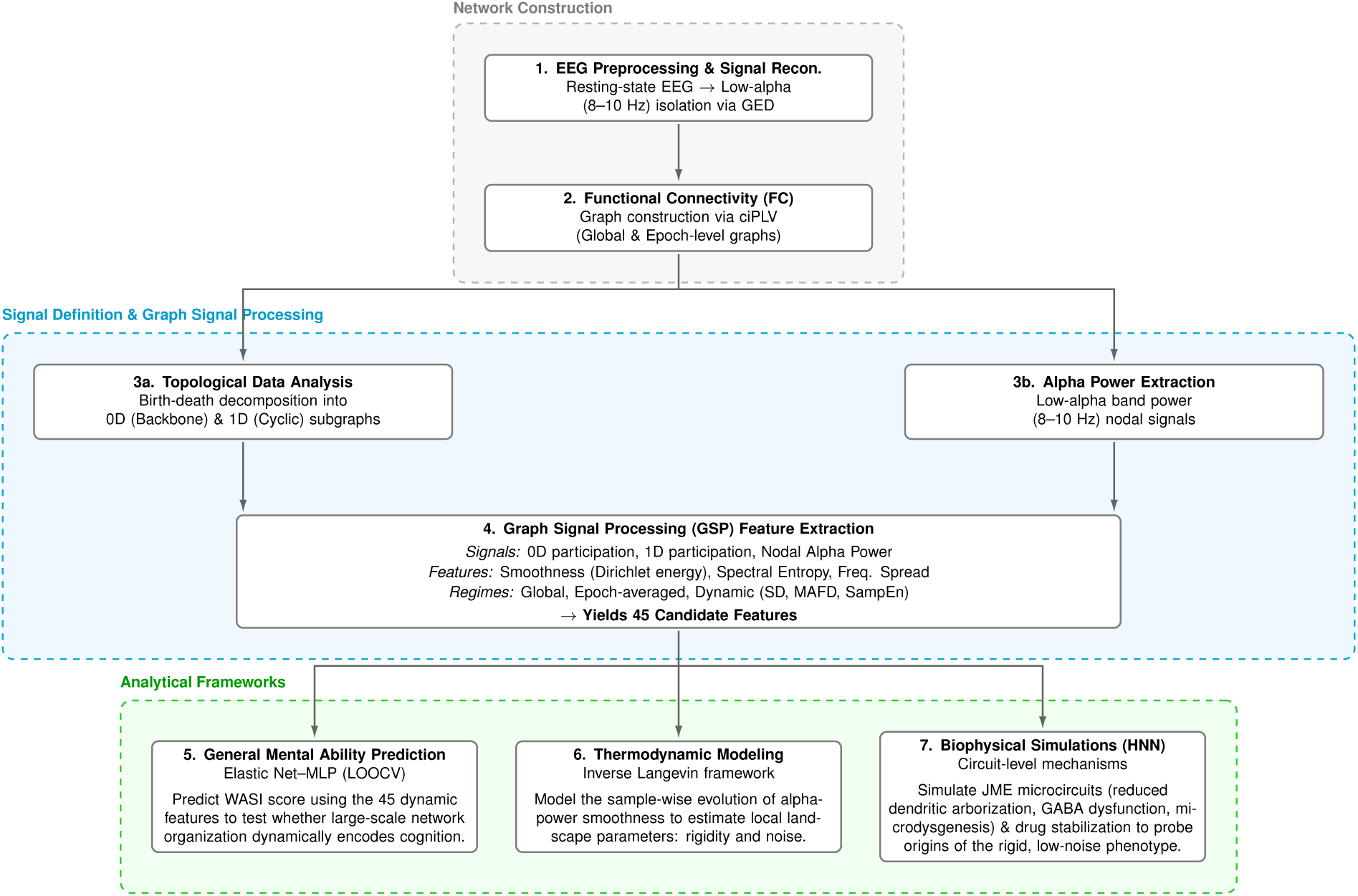
Overview of the analytical framework. Resting-state EEG was preprocessed and low-alpha activity was reconstructed using generalized eigendecomposition. Functional connectivity graphs were then computed and combined with topological data analysis and graph signal processing to derive graph-based descriptors from 0D participation, 1D participation, and nodal alpha-power signals across global, epoch-averaged, and dynamic regimes. These features were used to predict general mental ability, after which inverse Langevin modeling and biophysical simulations were used to characterize thermodynamic constraints and candidate circuit-level mechanisms.

In this framework, graph smoothness quantifies how gradually a signal varies across strongly connected parts of a network and is therefore linked to the expression of harmonic brain states on the functional connectome. We treated sample-wise alpha-power smoothness as a fluctuating state variable and used an inverse Langevin approach to reconstruct the effective landscape governing its temporal evolution, creating an interpretable low-dimensional description of a complex dynamical process. In this setting, thermodynamic rigidity refers to the degree to which ongoing fluctuations are confined around preferred smoothness states, with higher rigidity indicating less flexible exploration over time, whereas thermodynamic noise reflects the magnitude of local fluctuations around those states.

## Results

### General Mental Ability Prediction

We first asked whether EEG-derived graph features could predict individual differences in general mental ability, indexed by the raw estimated WASI full-scale score. In total, 45 candidate features were tested. These comprised three graph-signal descriptors (smoothness, spectral entropy, and frequency spread) computed on three nodal signal families—0D nodal participation (backbone-like structure), 1D nodal participation (cyclic structure), and nodal alpha power (AP)—in global and epoch-averaged regimes, together with dynamic summaries of the nine epoch-averaged features using standard deviation (STD), mean absolute first difference (MAFD), and sample entropy (SampEn). A hybrid Elastic Net–multilayer perceptron (MLP) model, evaluated using leave-one-out cross-validation (LOOCV), successfully predicted general mental ability in the control group, revealing a modest but statistically significant positive association between predicted and observed scores (Pearson *r* = 0.245, *p* = 0.0452; Fig. 2). By contrast, the same predictive pipeline failed in the JME group (*r* = −0.117, *p* = 0.6552), indicating that the EEG-derived feature space captured meaningful inter-individual variation in general mental ability in controls but not in patients.

**Figure 2:**
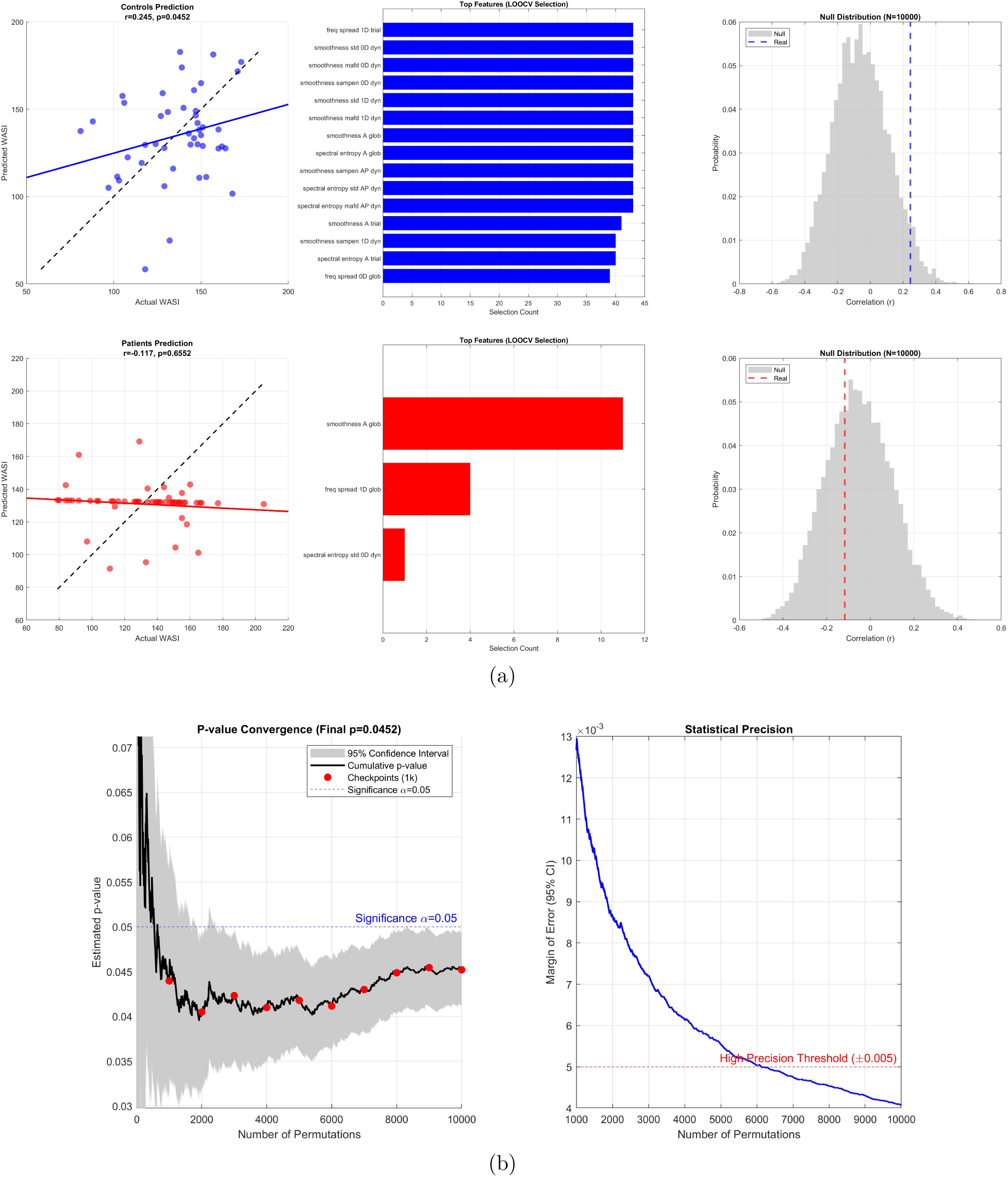
Prediction of general mental ability from EEG-derived network features and convergence of permutation-based significance testing. **a**, Predicted versus observed general mental ability scores, most frequently selected features across leave-one-out folds, and corresponding null distributions from 10,000 permutations for controls (top) and patients (bottom). **b**, Cumulative permutation-derived *p*-value estimate and 95% confidence interval for the control model, together with the margin of error across permutations.

Feature-selection frequencies further highlighted this group difference. In controls, the predictive signal was distributed across a stable subset of features: 14 of the 45 candidate features were selected in at least 40 of the 45 LOOCV folds. This stable subset was dominated by smoothness measures, particularly dynamic smoothness of the 0D and 1D participation signals, together with alpha-power smoothness and alpha-power spectral entropy features. This pattern indicates that general mental ability prediction in controls was driven primarily by temporal variation in how backbone-like, cyclic, and alpha-power nodal signals were organized on the same functional graph. In contrast, feature selection in patients was sparse and unstable. Only three features were selected at all across the entire LOOCV procedure, and the most frequently selected feature—global alpha smoothness—was selected in only 11 folds.

Permutation testing confirmed that the control-group result was unlikely under the null model, with the observed correlation falling in the extreme right tail of the null distribution generated from 10,000 label permutations (Fig. 2). The patient-group result was indistinguishable from the corresponding null distribution.

Because the control-group model was nominally significant, we additionally evaluated the stability of the permutation-based *p*-value estimate. The cumulative *p*-value trajectory stabilized after approximately 6,000 permutations, and the final estimate at 10,000 permutations was *p* = 0.0452, with an approximate 95% confidence interval of 0.041–0.049. The corresponding margin of error fell below the prespecified high-precision threshold of ±0.005, indicating that the permutation count was sufficient to support a stable inferential conclusion (Fig. 2).

### Thermodynamic Constraints on General Mental Ability

Because smoothness (Dirichlet energy) emerged as an important predictor in the control-group model, we next asked whether moment-to-moment fluctuations in alpha-power smoothness were related to general mental ability. This analysis focused on sample-wise alpha-power smoothness on the functional graph, because alpha power was the only signal available at the temporal resolution required for inverse Langevin modeling. The motivation for Langevin modeling is that it provides an interpretable low-dimensional description of a complex dynamical process by separating the tendency to return toward preferred states from the stochastic fluctuations around them. We estimated two local Langevin-derived parameters of this smoothness landscape near equilibrium: central rigidity, which quantifies how strongly the system is pulled back toward preferred smoothness states, and central noise, which quantifies the magnitude of local fluctuations around those states. For readability, we refer to these parameters below simply as rigidity and noise. We tested whether either parameter was associated with general mental ability.

In controls, rigidity was unrelated to general mental ability (robust regression slope *β* = −0.00, *p* = 0.992; Fig. 3). In patients, however, rigidity showed a significant inverse association with general mental ability (robust *β* = −0.12, *p* = 0.045), such that greater rigidity was associated with lower general mental ability.

**Figure 3:**
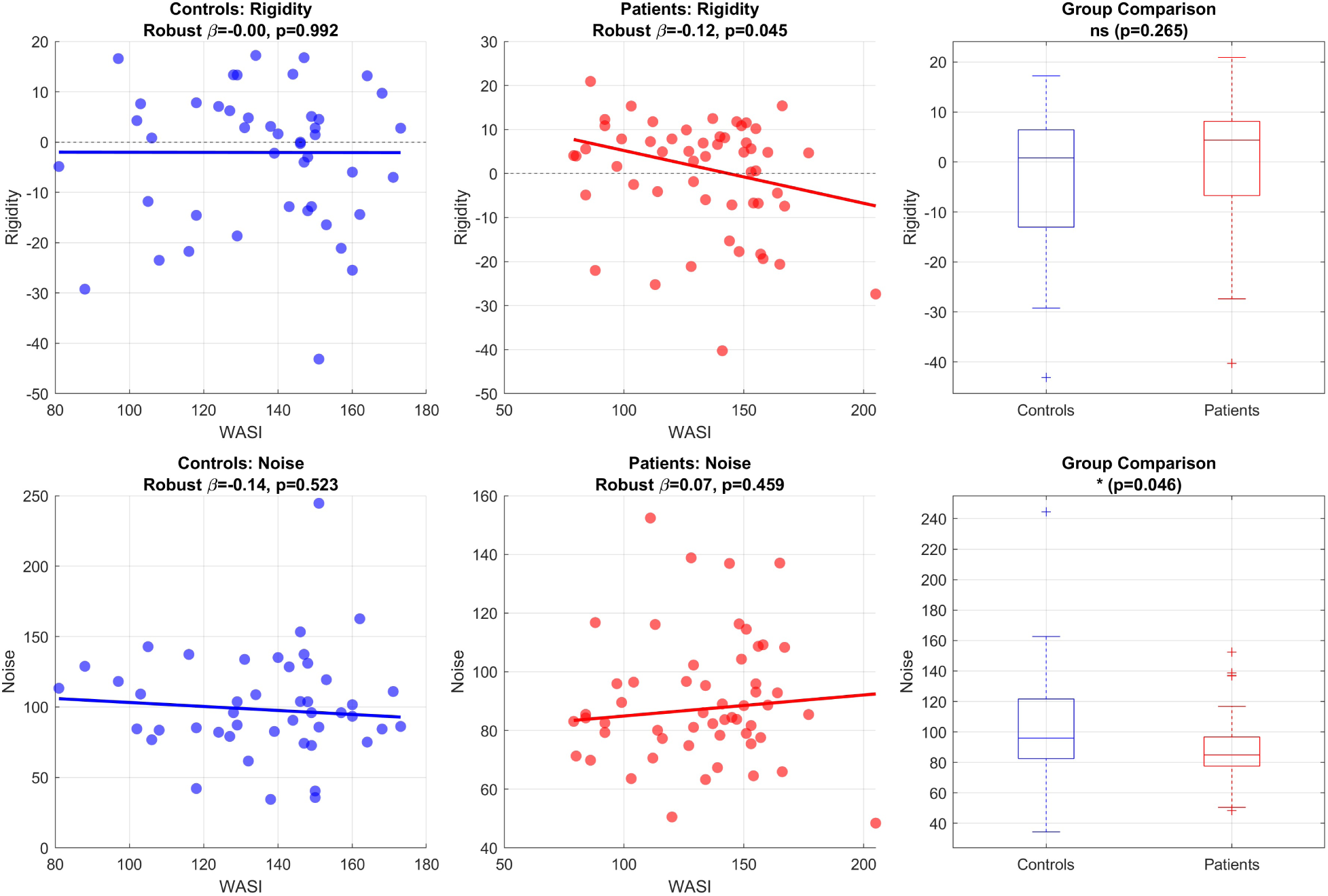
Thermodynamic parameters and general mental ability. Top row, associations between rigidity and general mental ability (WASI) in controls and patients, together with the between-group comparison of rigidity. Bottom row, analogous analyses for noise. Rigidity was inversely associated with general mental ability in patients but not controls, whereas noise was significantly lower in patients at the group level.

Rigidity did not differ significantly between groups (*p* = 0.265; Fig. 3). Thus, the key rigidity finding was not that patients, on average, showed greater rigidity than controls, but that within the patient group, individuals with greater rigidity were more likely to show lower general mental ability scores.

Noise showed a different pattern. Noise was not significantly associated with general mental ability in either controls (robust *β* = −0.14, *p* = 0.523) or patients (robust *β* = 0.07, *p* = 0.459). However, direct group comparison revealed significantly lower noise in patients than in controls (*p* = 0.046; Fig. 3), indicating that JME was characterized by weaker local fluctuations around equilibrium, even though inter-individual variation in noise did not itself predict general mental ability within either group.

### Biophysical Mechanisms of Thermodynamic Rigidity

To investigate the circuit-level mechanisms that could generate rigid smoothness landscapes, we used the Human Neocortical Neurosolver (HNN) (Neymotin et al., 2020) to simulate three candidate microcircuit alterations implicated in JME: reduced dendritic arborization, GABAergic dysfunction, and microdysgenesis. These simulations revealed a clear dissociation between mechanisms associated with hyperexcitability (GABAergic dysfunction and microdysgenesis) and the mechanism capable of preserving thermodynamic rigidity (reduced dendritic arborization) (Fig. 4).

**Figure 4:**
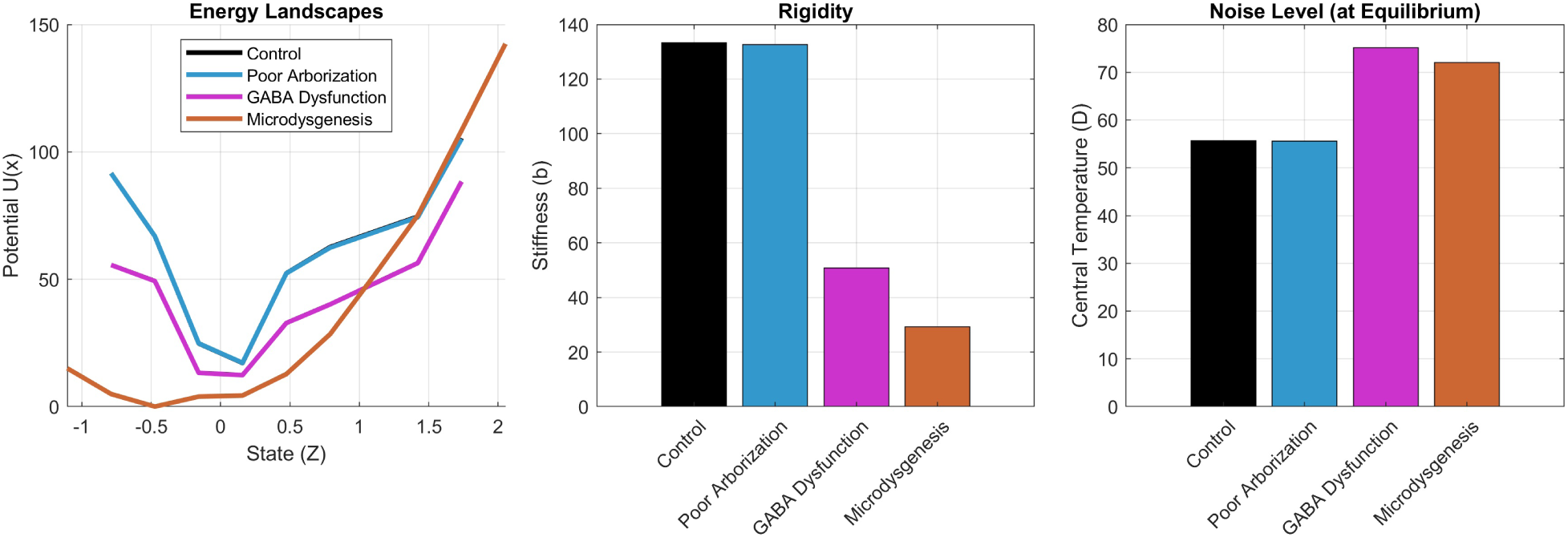
In-silico thermodynamic signatures of candidate JME microcircuit mechanisms. Left, reconstructed effective energy landscapes for the control, reduced dendritic arborization (poor arborization), GABA dysfunction, and microdysgenesis models. Middle, corresponding rigidity estimates. Right, central noise (central temperature) estimates. Poor arborization reproduced a high-rigidity, near-baseline-noise phenotype, whereas GABA dysfunction and microdysgenesis produced lower rigidity and elevated noise.

The reduced dendritic arborization model preserved a steep potential well and control-like rigidity, while maintaining near-control noise. In contrast, the GABA-dysfunction and microdysgenesis models produced shallower wells, substantially lower rigidity, and elevated noise relative to control. Thus, reduced dendritic arborization was the only pathological manipulation that maintained a rigid thermodynamic regime, whereas hyperexcitable circuit defects shifted the system toward a looser, more fluctuation-prone regime.

This dissociation suggests that thermodynamic rigidity is not a generic consequence of simulated pathology or hyperexcitability. Instead, it points more specifically to impaired dendritic integration as a plausible substrate of rigid smoothness dynamics. At the same time, because reduced dendritic arborization preserved near-control rather than reduced noise, it did not by itself reproduce the full thermodynamic pattern observed in patients, motivating the additional pharmacological simulations described below.

### Pharmacological Induction of Rigidity

We then tested whether simulated treatment with two commonly used anti-seizure medications, valproate (VPA) and levetiracetam (LEV), could shift these model-derived thermodynamic regimes. In the GABA-deficit model, VPA increased rigidity by 154% and LEV by 93%, with both treatments accompanied by clear reductions in noise (Fig. 5). In the microdysgenesis model, rigidity increased by 317% with VPA and by 193% with LEV. VPA also reduced noise substantially in this model, whereas LEV had only a modest effect on noise.

**Figure 5:**
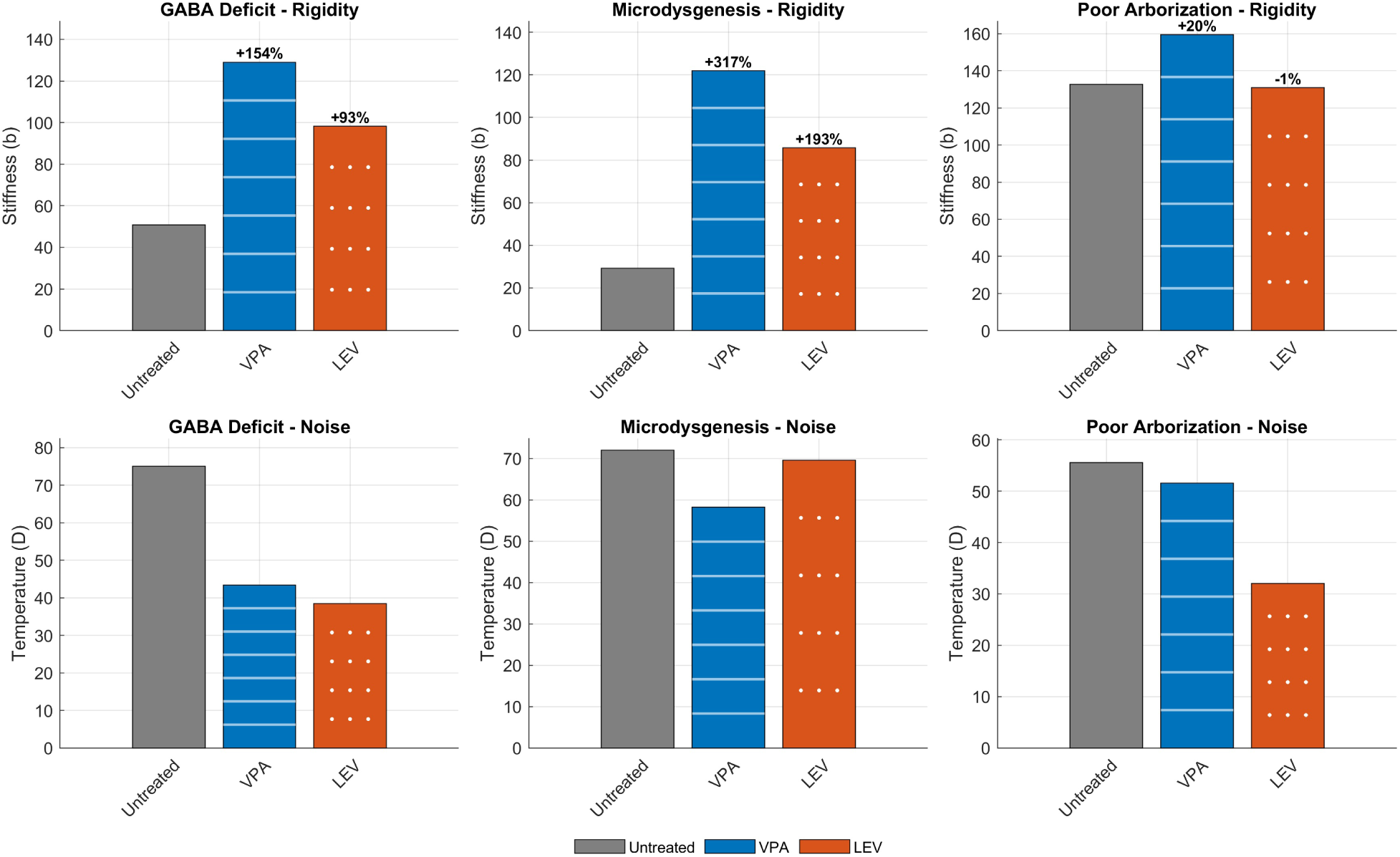
Effects of simulated pharmacological stabilization on thermodynamic parameters. Top row, changes in rigidity (stiffness) under untreated conditions and after simulated valproate (VPA) or levetiracetam (LEV) treatment in the GABA-deficit, microdysgenesis, and poor-arborization models. Bottom row, corresponding changes in noise (temperature). Pharmacological stabilization produced the largest rigidity increases in the hyperexcitable models, with more modest effects in the intrinsically rigid poor-arborization model.

In the poor-arborization model, which was already intrinsically rigid in the untreated state, treatment effects on rigidity were small. VPA produced only a modest additional increase in rigidity (+20%), whereas LEV left rigidity essentially unchanged (−1%). Noise was reduced only slightly by VPA but more clearly by LEV in this model.

Taken together, these simulations indicate that pharmacological stabilization most strongly reorganized the thermodynamic landscape in the hyperexcitable models, shifting them toward a more rigid and generally lower-noise regime. By contrast, when rigidity was already present in the untreated state, as in the poor-arborization model, treatment had only limited additional effects on rigidity.

## Discussion

In this study, we combined topological data analysis (TDA), graph signal processing (GSP), machine learning, inverse Langevin modeling, and biophysical simulation to examine how resting-state EEG network dynamics relate to general mental ability in juvenile myoclonic epilepsy (JME). Three principal findings emerged. First, general mental ability in healthy controls was modestly but significantly predicted by dynamic EEG-derived graph features, whereas the same predictive framework failed in JME. Second, within the patient group, greater thermodynamic rigidity was associated with lower general mental ability, and patients also exhibited lower thermodynamic noise than controls. Third, in silico modeling suggested that reduced dendritic arborization and pharmacological stabilization can both move the system toward a more rigid regime, although they do so from different mechanistic starting points.

### Dynamic Encoding of General Mental Ability in Health and Disease

The predictive results support the view that, in healthy individuals, general mental ability is encoded more strongly in dynamic network organization than in static summaries alone. In the control group, the most consistently selected features were dominated by temporal variability in topological smoothness, together with a smaller contribution from alpha-power smoothness and spectral entropy. This pattern suggests that general mental ability is related not simply to static summaries of alpha-related activity, but to how that activity is organized on the functional network scaffold over time.

This interpretation is broadly compatible with graph-spectral and harmonic accounts of large-scale brain dynamics, in which neural activity is represented as a time-varying mixture of graph modes ordered by smoothness or spatial scale (Atasoy et al., 2017; Glomb et al., 2020). In that sense, the harmonic brain states considered here can be understood as standing-wave-like patterns on the functional connectome whose relative expression changes over time. The present results suggest that, in healthy individuals, general mental ability is related to the flexibility with which activity is redistributed across these graph modes. This view is also consistent with broader accounts of metastable neural coordination, which emphasize that flexible cognition depends on controlled transitions among partially stable network states rather than persistence within a single optimal configuration (Tognoli & Kelso, 2014; Brinkman et al., 2022). Within this framework, the dominant role of dynamic smoothness features in controls suggests that general mental ability is supported by the ability to flexibly redistribute activity across the graph-defined backbone and cyclic subgraphs of the resting-state network.

By contrast, the same predictive framework failed in JME. This failure was not simply a reduction in effect size, but was accompanied by highly unstable feature selection, indicating that the model could not identify a coherent EEG-derived feature set that reliably mapped onto inter-individual variation in general mental ability in the patient group. One parsimonious interpretation is that JME preserves ongoing network fluctuations but disrupts their informative structure: the system still moves, but those movements are less consistently related to general mental ability. This distinction suggests that the core abnormality may lie in how network dynamics are organized, rather than merely in how much variability is present.

The prominence of low-alpha features in this analysis is also physiologically plausible. Alpha rhythms are strongly implicated in large-scale coordination, attentional control, and the temporal organization of cortical communication, making them a reasonable carrier frequency for the distributed dynamic effects interrogated here (Chapeton et al., 2019; Foster & Awh, 2019). Our findings therefore support the view that, in JME, vulnerability in general mental ability may be reflected less in static descriptors of alpha-related activity than in how alpha-band activity is moment-by-moment embedded in, and constrained by, the underlying functional network. More broadly, the present study evaluated only one biologically motivated configuration of this framework, namely low-alpha nodal signals analyzed relative to subject-specific low-alpha functional graphs. Other graph sources, frequency bands, and cross-frequency graph–signal combinations may capture additional variance relevant to general mental ability and may help clarify which aspects of large-scale brain dynamics are most informative for cognition.

### Thermodynamic Constraints and the Rigid-Brain Phenotype

To move from descriptive dynamics to an interpretable dynamical mechanism, we modeled the sample-wise temporal evolution of alpha-power smoothness in each subject using an inverse Langevin framework. This analysis revealed a dissociation between healthy individuals and patients. In controls, neither rigidity nor noise was significantly associated with general mental ability. In patients, however, rigidity showed a significant inverse relationship with general mental ability, such that greater confinement of the reconstructed smoothness landscape was associated with lower general mental ability.

This result sharpens the interpretation of the negative machine-learning finding in JME. The issue is not merely that general mental ability is less predictable in patients, but that a specific dynamical constraint—greater confinement around preferred smoothness states—appears to act as a cognitive bottleneck. Put differently, resting-state fluctuations remain present, but the accessible portion of the dynamic landscape becomes more tightly constrained in patients with lower general mental ability. In that sense, rigidity may not be a simple case-control biomarker, but rather a patient-specific limiting factor that becomes relevant only once the syndrome is present.

An important nuance is that rigidity did not differ significantly between groups at the population level. Thus, the key rigidity result is not a global upward shift in mean rigidity in JME, but a change in the functional meaning of rigidity: in controls, rigidity appears largely decoupled from general mental ability, whereas in patients it becomes behaviorally relevant. This argues against a simplistic interpretation in which JME is uniformly characterized by a more rigid landscape. Instead, the data are more consistent with a context-dependent or subgroup-dependent role for rigidity within the syndrome.

The noise result complements this picture. Noise did not significantly predict general mental ability within either group, but it was significantly lower in patients than in controls. We therefore interpret lower noise primarily as a group-level background condition rather than a direct determinant of general mental ability in this dataset. Conceptually, lower noise could reduce the probability of escaping local attractors and thereby amplify the behavioral consequences of rigidity, a view that is consistent with energy-landscape and stochastic-resonance accounts of brain dynamics (Watanabe et al., 2014; Kitajo et al., 2003; van der Groen et al., 2018; Nartallo-Kaluarachchi et al., 2026). At the same time, the absence of a significant within-group association between noise and general mental ability means that this interpretation should remain cautious. Our data support the presence of a colder (less noisy) equilibrium regime in JME, but they do not by themselves show that increasing noise would improve general mental ability.

### Biophysical Origins and a Two-Route Account of Rigidity

The biophysical simulation results help disentangle which circuit-level changes could plausibly generate the thermodynamic profile observed in the clinical data. The most notable finding was a dissociation between primary hyperexcitability and rigidity. In the untreated simulations, GABAergic dysfunction and microdysgenesis produced a relatively loose, high-noise regime, with reduced rigidity and increased noise. By contrast, reduced dendritic arborization produced a relatively rigid regime with near-baseline noise.

This distinction is important because it suggests that rigidity is not simply a generic consequence of epileptogenic microcircuit pathology. In our simulations, the models most directly associated with hyperexcitability—GABAergic dysfunction and microdysgenesis—did not reproduce the rigid phenotype linked to lower general mental ability in patients. Instead, the clearest rigid regime emerged in the reduced-dendritic-arborization model. One interpretation is that rigidity may arise more readily from impaired integrative capacity than from hyperexcitability alone.

Pharmacological stabilization further sharpened this picture. When valproate or levetiracetam was applied to the hyperexcitable models, rigidity increased sharply and noise generally decreased, shifting those models toward a more rigid and less noisy regime. In contrast, treatment had only modest additional effects on the already-rigid reduced-dendritic-arborization model. Taken together, these findings suggest a two-route account of rigidity in JME. One route is intrinsic, in which reduced dendritic arborization or structural disconnection promotes rigidity directly. The other is stabilizing, in which treatment suppresses noise and increases effective confinement in networks that would otherwise remain in a loose, hyperexcitable state.

This two-route account is necessarily speculative, but it offers a useful conceptual synthesis of the clinical and simulation results. It also fits with the broader pharmacological literature showing that anti-seizure medications stabilize neuronal excitability through mechanisms that can reduce effective network responsiveness and reactivity (Rogawski & Löscher, 2004). Within that framework, better seizure suppression does not necessarily imply greater cognitive flexibility: a treatment that is highly effective in preventing pathological excursions may also reshape the resting-state landscape in a way that is less favorable for flexible cognitive dynamics. Importantly, this interpretation does not imply that anti-seizure medications are the sole cause of cognitive difficulty in JME, but rather that treatment-related stabilization may contribute to a cognitively less flexible dynamical regime in a subset of patients.

### Relationship to Prior EEG Evidence in JME

Routine EEG background is often described as normal in JME, but quantitative EEG studies have long suggested that more subtle abnormalities are present. Prior work has reported altered spectral power, coherence, and background organization in JME, even in recordings that appear visually unremarkable on routine inspection (Santiago-Rodríguez et al., 2008; Tikka et al., 2013; Santiago-Rodríguez et al., 2022). Our findings extend that literature in two ways.

First, they suggest that abnormalities relevant to general mental ability may be better captured in the embedding of oscillatory activity within network structure than in static descriptors alone. In our data, the most informative features in healthy individuals were dynamic graph-derived descriptors, especially topological smoothness, rather than static graph-based summaries. Second, the results suggest that the pathophysiology relevant to general mental ability in JME may lie not only in altered oscillatory content, but in altered state-space organization: how easily the system moves among graph-constrained activity patterns over time.

This perspective also aligns with the broader contemporary view of JME as a network disorder with distributed frontal-thalamocortical involvement rather than a narrowly defined seizure syndrome (Anderson & Hamandi, 2011; Wandschneider et al., 2012; Garcia-Ramos et al., 2025). In that sense, the present framework may be useful because it links three levels of description that are often studied separately: oscillatory physiology, large-scale network topology, and cognitively relevant dynamical constraints.

### Limitations and Future Directions

Several limitations should be acknowledged. First, the predictive effect in controls was modest and the machine-learning analysis was performed in a relatively small sample without external validation. The control-group result should therefore be interpreted as proof of principle rather than as a finalized biomarker. In held-out data, the observed correlation corresponded to approximately 6% shared variance between predicted and observed scores (*r*^2^ ≈ 0.06). We therefore do not interpret the present framework as capturing a dominant determinant of general mental ability, but rather as evidence that graph-spectral organization of resting-state EEG contains behaviorally meaningful information despite sampling only a small portion of the possible graph–signal design space.

Second, the study is cross-sectional, which limits causal inference. In particular, the clinical data cannot by themselves disentangle the relative contributions of syndrome biology, seizure burden, medication exposure, and developmental history to the observed thermodynamic signatures. An additional methodological consideration is that the functional graph and several of the analyzed nodal signals were derived from the same subject-specific electrophysiological dataset. This is standard in graph-spectral analyses, where a graph provides the geometry relative to which a nodal signal is evaluated, but it also means that measured smoothness reflects signal organization relative to a geometry inferred from related data rather than from an entirely independent scaffold. Future work should test how strongly the present findings depend on this design choice by comparing alternative graph constructions, including structural graphs, cross-frequency functional graphs, and other graph–signal pairings.

Third, the thermodynamic analysis relies on a coarse-grained Langevin approximation. Although this framework is useful for summarizing state-dependent drift and diffusion, it remains an approximation to high-dimensional neural dynamics and should not be interpreted as a literal mechanistic model of the brain. Fourth, the HNN simulations are intentionally simplified. They model a canonical cortical column rather than a full thalamocortical or whole-brain system, and the medication manipulations are phenomenological rather than pharmacokinetic. The simulations should therefore be interpreted as hypothesis-generating rather than definitive evidence for any specific cellular mechanism.

Fifth, several key inferential results were modest and fell near the conventional 0.05 threshold. Although the thermodynamic analyses were targeted rather than broad feature-screening tests, these findings should be interpreted cautiously until replicated in independent cohorts. Sixth, the cognitive outcome was the raw estimated WASI full-scale score rather than the age-standardized index, because the aim was to preserve developmentally meaningful variance that may be reflected in brain-network organization. Accordingly, the observed associations should be interpreted as relating to general mental ability across the sampled age range, rather than to intellectual performance normalized relative to age peers.

These limitations point directly to future work. External validation in independent cohorts will be required to determine whether the predictive and thermodynamic findings generalize. Longitudinal studies incorporating medication changes, seizure control, and repeated cognitive testing will be important for distinguishing intrinsic rigidity from treatment-related rigidity. It will also be important to test whether the present framework extends beyond JME to other generalized and focal epilepsies, and whether thermodynamic descriptors can identify patient subgroups in whom cognitive risk is disproportionately linked to network confinement—whether intrinsic, treatment-related, or both—rather than to ongoing epileptic burden alone. Finally, a structured comparison of graph–signal pairings, including same-band and cross-band combinations, will be needed to determine whether the present low-alpha configuration is uniquely informative or simply one informative point within a broader harmonic description of cognition-relevant brain dynamics.

## Conclusion

In summary, the present findings support a model in which general mental ability in healthy individuals is reflected in flexible, dynamically informative resting-state network fluctuations, whereas in JME this relationship becomes degraded and, in some patients, constrained by a more rigid thermodynamic landscape. The key implication is not simply that patients fluctuate differently, but that the meaning of those fluctuations changes: in JME, greater rigidity appears to become a cognitive liability. The simulation results further suggest that this liability may arise through more than one mechanistic route, including both intrinsic reductions in dendritic integration and treatment-related stabilization of otherwise hyperexcitable networks.

Viewed in this way, cognitive dysfunction in JME may reflect not only abnormalities of excitation and inhibition, but also abnormalities in how the brain explores its accessible dynamic landscape. This framework does not diminish the importance of seizure control. Rather, it suggests that the long-term optimization of clinical care may eventually require balancing seizure suppression with preservation of sufficient dynamical flexibility to support cognition.

## Methods

The institutional review board of the University of Wisconsin (Health Sciences IRB, ID 2019-1416) approved all study procedures. All participants provided written informed consent. Procedures complied with relevant guidelines and regulations. All data were de-identified prior to analysis, and no protected health information was included in the analytic dataset.

### Participants

We studied 99 individuals from the Comprehensive Epilepsy Center of the University of Wisconsin Department of Neurology with high-quality resting-state EEG recordings: 45 healthy controls (19 male, 26 female; mean age 18.8 years) and 54 patients with juvenile myoclonic epilepsy (JME) (16 male, 38 female; mean age 19.3 years). The full cohort had a mean age of 19.0 years and included 35 males and 64 females.

JME participants were eligible if they were 12–25 years old and met diagnostic criteria for JME, including myoclonic jerks, at least one afebrile generalized tonic–clonic seizure not attributable to metabolic disturbance or acute traumatic injury, and EEG findings of generalized 3.5–5 Hz spike-wave and/or polyspike discharges (Baykan & Wolf, 2017). Exclusion criteria were recent or remote infarction, neoplasm, or cerebral dysgenesis; other significant neurologic or neurodevelopmental disorders; English as a second language; or verbal or performance IQ *<* 70.

Healthy controls were recruited through community outreach and related epilepsy research studies. Eligible controls were 12–25 years old at enrollment and were either unrelated community volunteers or biological relatives (siblings or first-degree cousins), or friends/significant others, of individuals with epilepsy. Exclusion criteria were English as a second language; any initial precipitating event (e.g., febrile seizures); any history of seizure-like episodes; or any diagnosed neurologic or psychiatric disorder; or verbal or performance IQ *<* 70.

Initial EEG quality screening was based on visual inspection of the resting-state recordings. A second quality-control step was applied after generalized eigendecomposition (GED)-based alpha reconstruction, and subjects whose reconstructed low-alpha activity did not show the expected posterior-dominant scalp distribution were excluded from further analysis.

At the time of EEG acquisition, all patients with JME were receiving anti-seizure medication under usual clinical care. The mean number of anti-seizure medications per patient was 1.48. The most commonly used medications were levetiracetam (57%), lamotrigine (31%), and valproate (18%).

### Cognitive Assessment

General mental ability was indexed using the 4-subtest Wechsler Abbreviated Scale of Intelligence (WASI), comprising Vocabulary, Similarities, Block Design, and Matrix Reasoning (Wechsler, 1999). The dependent measure was the raw estimated Full Scale IQ score (IQFULLR), rather than the age-standardized index, because the goal was to preserve developmentally meaningful variance across the sampled age range. In this study, age-related variation was therefore treated as part of the neurodevelopmentally relevant construct of interest rather than as a nuisance factor to be regressed out.

### EEG Acquisition

High-density EEG was acquired using a 256-channel cap (Quick-Cap Neo Net, Compumedics) connected to a Compumedics Neuroscan system with four Neuvo 64-channel amplifiers at 1000 Hz. For each subject, 5–10 minutes of eyes-closed, awake resting-state data were collected.

### Preprocessing

Continuous resting-state EEG was preprocessed using the RELAX automated pipeline (v2.0.0) implemented in EEGLAB (Delorme & Makeig, 2004; Bailey et al., 2023a; Bailey et al., 2023b; Bailey et al., 2025). Non-scalp electrodes were removed. Data were filtered with a 0.5 Hz high-pass filter, a 100 Hz low-pass filter, and a 60 Hz notch filter. Noisy scalp channels were identified using the PREP pipeline (Bigdely-Shamlo et al., 2015). Artifact attenuation was performed using a multi-pass multi-channel Wiener filter, followed by independent component analysis using PICARD (Somers et al., 2018; Ablin et al., 2018). Independent components classified by ICLabel as eye- or muscle-related with probability *>* 0.80 were cleaned using targeted wavelet-enhanced ICA, after which the data were average-referenced and previously rejected scalp channels were interpolated (Pion-Tonachini et al., 2019). For downstream analyses, data were resampled to 100 Hz using EEGLAB’s `pop_resample`, which applies anti-aliasing filtering during resampling.

### Signal Reconstruction via Generalized Eigendecomposition

A fixed 180-second segment was analyzed for all subjects to ensure a common number of epochs and directly comparable dynamic and thermodynamic estimates across participants despite variability in total recording duration and usable data length. The first 15 seconds were discarded because the beginning of resting-state recordings is commonly more susceptible to settling and movement-related contamination. The subsequent 180 seconds were segmented into 60 non-overlapping 3-second epochs. This epoch length was chosen as a compromise between reliable alpha-band functional-connectivity estimation and sufficient temporal resolution for dynamic analyses.

Generalized eigendecomposition (GED) was used to isolate subject-specific low-alpha activity (Cohen, 2022). For each subject, a signal covariance matrix (*S*) was computed from 8–10 Hz filtered data and a reference covariance matrix (*R*) from broadband data. Both matrices were mean-centered and regularized with Ledoit–Wolf shrinkage (Ledoit & Wolf, 2004). The generalized eigenvalue problem was solved as

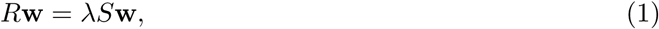

where **w** denotes a spatial filter and *λ* the associated eigenvalue. This formulation is algebraically equivalent to the more common *S***w** = *λR***w** form, but yields eigenvalues that reflect the ratio of broadband to low-alpha variance. Components were therefore sorted in ascending order, so smaller eigenvalues corresponded to stronger low-alpha enrichment relative to the broadband reference.

The number of retained components was determined within a bounded range of 3–12 using a relative stability criterion. Components were retained up to the first index at which the difference between consecutive eigenvalues dropped below 5% of the preceding eigenvalue, with a minimum of 3 and a maximum of 12 components. This bounded criterion was used to avoid retaining trivial low-variance components while allowing subject-specific variation in the dimensionality of the low-alpha subspace.

Component time series were obtained by projecting the retained spatial filters onto the mean-centered 8–10 Hz data. For each retained component, a spatial map was estimated from the alpha covariance structure, and polarity was corrected by identifying the channel with the largest absolute loading and flipping both the spatial map and time series if that loading was negative. Spatial maps were then min–max normalized, i.e., linearly rescaled to the interval [0, 1], to place component weights on a common nonnegative scale before back-projection. The final electrode-level alpha time series was reconstructed by weighted back-projection of the retained components, with each component weighted by the inverse of its generalized eigenvalue so that components with stronger alpha-to-broadband separation contributed more strongly to the final signal (Cohen, 2022).

### Functional Connectivity Graph Calculation

Functional connectivity (FC) graphs were computed from the reconstructed alpha-band time series using the corrected imaginary phase-locking value (ciPLV) (Bruña et al., 2018).

#### Epoch-level graphs

For each 3-second epoch, ciPLV was computed using a windowed-average approach to reduce within-epoch non-stationarity. Each epoch was divided into 10 non-overlapping windows; complex-valued phase-locking value matrices were computed within each window and averaged before conversion to ciPLV. This yielded 60 epoch-specific FC matrices per subject.

#### Global graphs

A subject-level FC graph was obtained by first computing the complex-valued phase-locking value separately for each epoch, averaging across epochs, and then converting the epoch-averaged complex PLV to ciPLV.

All subsequent analyses used the absolute value of the resulting ciPLV matrices. This step yielded unsigned edge weights appropriate for graph-topological and graph-spectral analysis, because for the present purposes the sign of the phase relation was not interpreted as a separate graph feature; instead, we focused on connection magnitude when constructing FC graphs, graph Laplacians, and topological subgraphs.

### Defining Signals via Topological Data Analysis

Birth–death decomposition was applied to the absolute-valued ciPLV graphs for both the epoch-specific and global FC matrices (Chung et al., 2023). Here, topological data analysis (TDA) was used not as a standalone inferential endpoint, but as a principled way to decompose each FC graph into complementary subgraphs with distinct organizational meaning. Specifically, the decomposition separates the minimal set of edges required to maintain connectedness from the remaining redundant or cyclic edges.

The maximum spanning tree (MST) was identified as the 0D topological backbone, corresponding to the set of edges required to connect all nodes into a single component. In the language of persistent homology, these edges correspond to the 0D structure that supports graph connectedness. All remaining edges were assigned to the 1D topology, representing cyclic or redundant connectivity beyond the backbone (Chung et al., 2023).

For each subgraph, nodal signals were defined as node strength at each electrode and z-scored across electrodes. The resulting 0D signal therefore quantified each electrode’s participation in the backbone-like connectivity structure, whereas the 1D signal quantified participation in cyclic or redundant connectivity. In addition, low-alpha band power (8–10 Hz) of the reconstructed signal was computed for each electrode and epoch, yielding an alpha-power nodal signal complementary to the two topology-derived signals.

### Analyzing Signals via Graph Signal Processing

Graph signal processing (GSP) was used to quantify how these nodal signals were organized relative to the FC graph (Ortega et al., 2018). Analyses were performed in two settings: a static/global setting using global signals and global graphs, and a dynamic/epoch-based setting using epoch-specific signals and graphs.

For each graph, the normalized graph Laplacian was computed as

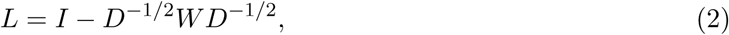

where *W* is the weighted adjacency matrix and *D* the diagonal degree matrix. Eigendecomposition of *L* yielded the graph harmonics (eigenvectors, collected in *U*) and graph frequencies (eigenvalues). Lower graph frequencies correspond to smoother, more spatially extended patterns on the graph, whereas higher frequencies correspond to sharper changes across connected nodes. For a nodal signal *x*, the graph Fourier transform was

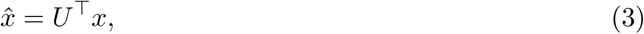

and squared coefficient magnitudes defined graph-spectral energy.

### Calculated Features and Analysis Regimes

Features were derived from three nodal signal families: the 0D topology signal, the 1D topology signal, and the alpha-power signal.

#### GSP features

For each signal, we computed three complementary graph-spectral descriptors. Smoothness was defined as

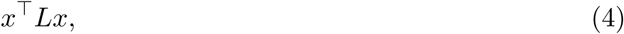

which corresponds to the graph Dirichlet energy and quantifies the extent to which a signal varies gradually across strongly connected parts of the graph (Ortega et al., 2018). Lower values indicate smoother signals, that is, signals whose energy is concentrated relatively more in lower graph frequencies.

Spectral entropy was computed as the Shannon entropy of the normalized graph-spectral energy distribution using the natural logarithm,

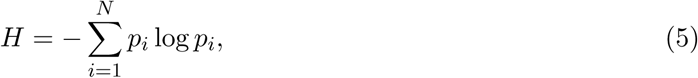

Where

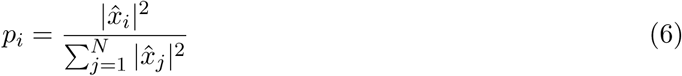

is the normalized energy at graph frequency *i*. This quantity captures how broadly signal energy is distributed across graph modes.

Frequency spread was computed as the energy-weighted standard deviation of graph frequencies,

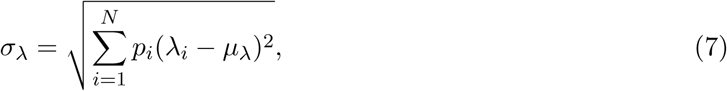

Where

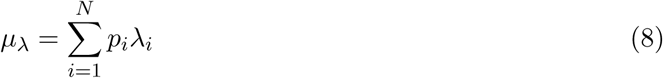

is the energy-weighted mean graph frequency. This quantity captures the dispersion of graph-spectral energy around its dominant graph scale.

#### Analysis regimes

Features were computed in three regimes. In the static/global regime, each feature was computed once per subject from the global signal and global graph. In the static/epoch-averaged regime, features were computed for each epoch and then averaged across the 60 epochs. In the dynamic regime, epochwise feature values were treated as a 60-point time series.

#### Dynamic features

For each dynamic feature time series, we computed the standard deviation (SD), mean absolute first difference (MAFD), and sample entropy (SampEn) (Richman & Moorman, 2000). SD captured overall between-epoch variability, MAFD captured step-to-step variability across consecutive epochs, and SampEn captured temporal irregularity or unpredictability. SampEn was computed with embedding dimension *m* = 2. The tolerance parameter *r* was defined separately for each signal family and feature type as 0.2× the median across-subject standard deviation of the corresponding feature time series. This scaling was chosen to place the tolerance on a comparable relative scale across feature families while remaining independent of the behavioral outcome.

In total, this yielded 45 candidate features per subject: 9 static/global features, 9 static/epoch-averaged features, and 27 dynamic summary features.

### Machine Learning and Statistical Validation

To predict general mental ability, we used a hybrid Elastic Net–multilayer perceptron (MLP) framework with leave-one-out cross-validation (LOOCV) (Wechsler, 1999; Zou & Hastie, 2005). The rationale for this two-stage approach was that sparse linear feature selection can identify a stable subset of informative predictors, whereas a shallow neural network can then model potentially nonlinear combinations among those retained features.

Within each LOOCV training fold, Elastic Net performed feature selection using *α* = 0.5, 25 candidate *λ* values, and internal 5-fold cross-validation to identify the minimum-MSE solution. Elastic Net was implemented using MATLAB’s `lasso` function with default predictor standardization. A shallow feedforward MLP with one hidden layer of 10 neurons was then trained on the selected features to predict the left-out subject’s raw estimated WASI full-scale score (IQFULLR). The network was trained using scaled conjugate gradient backpropagation (`trainscg`). Ten hidden units were chosen a priori as a simple low-capacity nonlinear model rather than tuned to maximize performance. A fixed random seed was used to ensure reproducibility of model initialization and training. If Elastic Net selected no features in a given training fold, the left-out subject was assigned the training-set mean IQFULLR.

Before model fitting, missing feature values were imputed with the corresponding column mean computed within the analyzed group. Model performance was quantified as the Pearson correlation (*r*) between observed and predicted scores. Significance was assessed by repeating the full LOOCV pipeline 10,000 times with randomly shuffled IQFULLR values to generate a null distribution. Feature importance was defined as feature-selection frequency across LOOCV folds. Univariate group comparisons and feature–behavior associations were assessed using non-parametric permutation testing with max-statistic control of the family-wise error rate (Nichols & Holmes, 2002).

To assess the stability of permutation-based inference, we tracked the cumulative *p*-value estimate across permutations. At iteration *N*, the cumulative *p*-value was

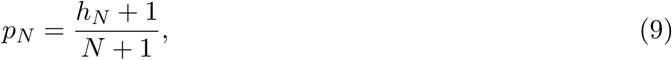

where *h_N_* is the cumulative number of null correlations greater than or equal to the observed correlation. The corresponding standard error was

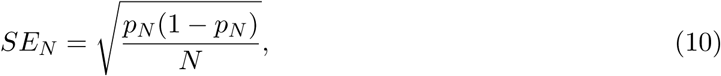

and the 95% Wald confidence interval was computed as *p_N_* ± 1.96 × *SE_N_*. Convergence was defined as a final margin of error below 0.005.

### Thermodynamic Modeling of Network Dynamics

To derive an interpretable low-dimensional description of a high-dimensional dynamical process, we modeled the temporal evolution of alpha-power smoothness using an inverse Langevin framework based on stochastic thermodynamics and the Kramers–Moyal expansion (Risken, 1996; Gardiner, 2009). In generic form, the dynamics of a one-dimensional state variable *z*(*t*) can be written as

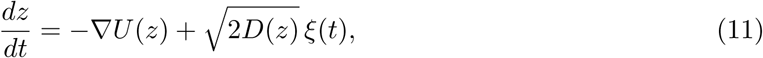

where −∇*U* (*z*) is the deterministic drift term, *D*(*z*) is the state-dependent diffusion term, and *ξ*(*t*) is Gaussian white noise. In this study, the goal was not to fit a mechanistically complete neural model, but to obtain empirical estimates of restoring tendency and fluctuation strength for a graph-derived brain-state variable.

#### State variable definition

Alpha power was defined as the squared amplitude of the reconstructed low-alpha signal. Alpha-power smoothness was then computed sample-by-sample as

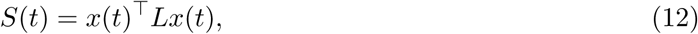

where *x*(*t*) is the vector of electrode-wise alpha power at time *t* and *L* is the normalized Laplacian of the epoch-specific FC graph corresponding to the epoch containing time point *t*. This quantity measures how gradually the spatial distribution of alpha power varies across strongly connected parts of the graph. Lower values indicate a smoother spatial pattern of alpha power across the functional network, whereas higher values indicate sharper spatial variation. Thus, the thermodynamic analysis was performed on the smoothness of alpha power rather than on the raw oscillatory signal itself.

#### Drift–diffusion estimation

For each subject, the sample-wise alpha-power smoothness time series was concatenated across epochs and z-scored globally. State increments were computed as Δ*z_t_* = *z_t_*_+Δ_*_t_* − *z_t_*, with Δ*t* = 1*/*100 s after resampling to 100 Hz. The state space was restricted to the central 95% of the empirical distribution (2.5th–97.5th percentiles) and divided into 20 linearly spaced bins. Twenty bins were used as a compromise between state-space resolution and sufficient sample occupancy per bin for stable drift–diffusion estimation. Within each state-space bin *k*, empirical drift and diffusion were estimated from the increments, using only bins containing more than 10 samples, as

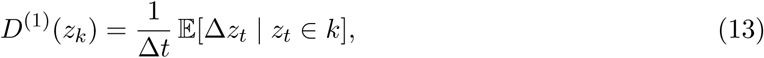

and

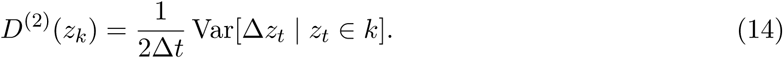

Subjects with fewer than 1000 valid smoothness samples after NaN removal were excluded, although no subject met this exclusion criterion. This procedure provides a coarse-grained empirical approximation of the deterministic drift and stochastic diffusion terms of the observed dynamics.

### Landscape Reconstruction and Metric Extraction

For each subject, the drift field was approximated by fitting a third-degree polynomial to the valid binwise drift estimates,

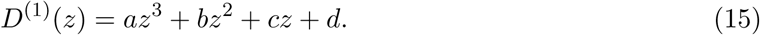

A cubic polynomial was used as the lowest-order flexible form capable of capturing both approximately harmonic single-well dynamics and asymmetric or bistable departures from that regime. The effective potential was then reconstructed from the drift by integrating its negative,

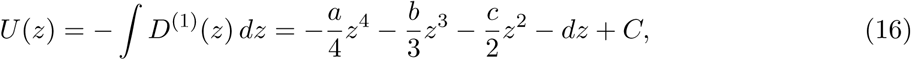

consistent with the Langevin relation between drift and the gradient of the potential landscape. Fits were retained only when at least four valid bins were available; all subjects met this criterion.

#### Thermodynamic metrics

Because the smoothness series was z-scored, thermodynamic metrics were defined relative to the standardized central operating regime, i.e., the neighborhood of *z* ≈ 0, rather than to a separately estimated physiological fixed point. Because rigidity was intended to summarize local confinement around this central regime, central rigidity was defined as the coefficient of the quadratic term in the reconstructed potential,

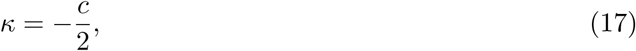

where *c* is the linear coefficient of the fitted drift polynomial. Near *z* = 0, this quadratic term provides the leading-order description of local confinement, whereas the cubic and quartic terms capture asymmetry and nonlinear structure farther from the center. Larger *κ* values therefore indicate stronger local confinement of fluctuations around the standardized central operating regime.

Central noise was defined as the mean diffusion coefficient within the interval *z* ∈ [−1, 1], which captures the magnitude of local fluctuations in the same central regime. For readability, these quantities were referred to in the Results and Discussion simply as rigidity and noise.

Associations between thermodynamic parameters and general mental ability were assessed separately in controls and patients using MATLAB’s `robustfit`, which implements iteratively reweighted least squares with the default bisquare weighting function. The predictor was the raw estimated WASI full-scale score (IQFULLR), and the outcome was either central rigidity or central noise. Reported robust *β* values correspond to the estimated slope coefficients, and associated *p*-values test the null hypothesis that the slope equals zero.

### In-Silico Biophysical Modeling of JME Pathophysiology

To examine candidate circuit mechanisms for the observed thermodynamic phenotype, we used the Human Neocortical Neurosolver (HNN) (Neymotin et al., 2020). Simulations were based on the Jones 2009 cortical column model, comprising layer 2/3 and layer 5 pyramidal neurons and inhibitory interneurons (Jones et al., 2009). Resting-state-like dynamics were generated by replacing the standard evoked drives with a continuous 10 Hz proximal burst drive and background distal Poisson input.

Three pathological model variants were tested. Reduced dendritic arborization was simulated as a 40% reduction in distal feedforward input strength, as a proxy for reduced dendritic spine density and branching complexity (Wong, 2005). GABAergic dysfunction was simulated as a 40% reduction in inhibitory synaptic weight from basket cells to pyramidal somata, approximating impaired *GABRA1*-related inhibition (Cossette et al., 2002). Microdysgenesis was simulated as a 40% increase in recurrent excitation among pyramidal neurons, modeling aberrant local clustering and cortical layering abnormalities described in primary generalized epilepsy (Meencke & Janz, 1984). Perturbation magnitudes were chosen to reflect partial rather than complete dysfunction, guided by the referenced experimental and neuropathological literature.

For each condition, 30 seconds of activity were simulated and somatic membrane voltages were extracted from all modeled cells. Signals were downsampled to 100 Hz and bandpass filtered to 8–10 Hz. Each condition was treated as a representative 30-second steady-state realization under fixed model parameters, chosen to characterize the thermodynamic regime associated with that parameter set rather than trial-to-trial variability across repeated stochastic runs. Because these simulated signals were recorded at the level of individual model cells and were not affected by volume conduction, functional connectivity was estimated using absolute Pearson correlation rather than ciPLV. The resulting correlation matrix was used as the graph for subsequent graph-signal and Langevin analyses.

### Pharmacological Validation

To test the thermodynamic consequences of anti-seizure treatment, we simulated valproate (VPA) and levetiracetam (LEV). VPA was modeled as a 50% increase in inhibitory synaptic weight. LEV was modeled as a 20% reduction in global synaptic gain, consistent with its SV2A-related reduction of neurotransmitter release probability (Lynch et al., 2004; Meehan et al., 2012). Treated models were analyzed using the same graph-signal and inverse Langevin framework applied to the untreated simulations.

## Data Availability

The raw data supporting this study are not publicly available because they contain protected health information and are subject to HIPAA and institutional privacy restrictions. De-identified derived features may be made available from the corresponding author on reasonable request, subject to institutional approval.

## Code Availability

The code used for data preprocessing, feature extraction, statistical analysis, and in silico modeling is publicly available at https://github.com/felipebpaiva/JMECP-EEG/tree/main.

## Acknowledgements

We thank Klevest Gjini and Dace Almane for their contributions to data collection and organization.

## Author Contributions

Felipe Branco De Paiva conceived the study, performed the analyses, acquired funding, and wrote and revised the manuscript. Aaron F. Struck supervised the study, acquired funding, and revised the manuscript. Moo K. Chung supervised the study and revised the manuscript. Bruce P. Hermann revised the manuscript. Meixian Zhao, Meishu Zhao, and Steven E. Haworth contributed to the analyses. Santiago Philibert-Rosas and Cameron J. Brace contributed to writing and revising the manuscript. Erika Moe contributed to writing the manuscript.

